# Discovery of New Classes of Glycine transporter 2 (GlyT2) Inhibitors and Study of GlyT2 Selectivity by Combination of Novel Structural Based Virtual Screening Approach and Free Energy Perturbation (FEP+) Calculations

**DOI:** 10.1101/510487

**Authors:** Filip Fratev, Manuel Miranda-Arango, Elvia Padilla, Suman Sirimulla

## Abstract

In recent years, the mammalian GlyT2 transporter has emerged as a promising target for the development of anti-chronic pain agents. In our current work, we discovered a new set of promising hits that inhibit the glycine transport at nano and micromolar activity and have excellent selectivity over GlyT1 (as shown by in vitro studies), using a newly designed virtual screening (VS) protocol that combines a structure-based pharmacophore and docking screens. Furthermore, the free energy perturbation (FEP+ protocol) calculations and molecular dynamics (MD) studies revealed the GlyT2 amino acid residues critical for the binding and selectivity of both Glycine and our Lead1 compound. The FEP+ results well-matched available literature mutational data proving the quality of the generated GlyT2 structure. Based on these calculations we propose that Lead1 may also be a strong inhibitor of the neutral and basic amino acid transporter B (0+) (SLC6A14). Thus, the subsequent lead optimization and characterization of refined compounds may lead to both chronic pain and pancreatic cancer agents addressing an unmet and challenging clinical needs.

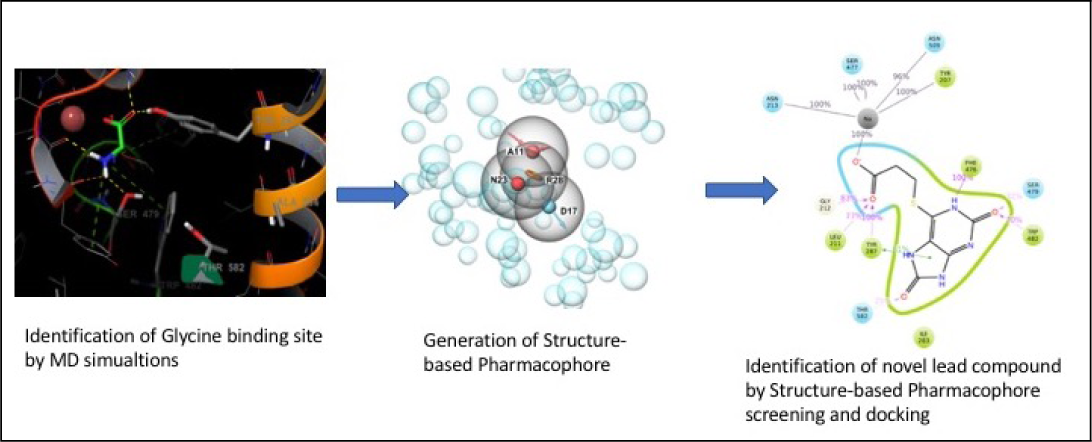

## Introduction

In the mammalian CNS, glycine and GABA function as inhibitory neurotransmitters by binding to postsynaptic chloride receptors, resulting in chloride influx and hyperpolarization. These glycinergic neurons participate in inhibition of pain signal transmission by releasing glycine onto pain fibers and dysfunction of these neurons leads to augmented excitation and transmission of neuropathic pain signals in the dorsal horn.^1,2^ The glycine transporter 2 (GlyT2) is responsible for glycine reuptake from the extracellular space back into the glycinergic neuron to terminate neurotransmission; therefore, pharmacological GlyT2 inhibition would allow prolonged glycine binding and activation of glyine receptor (GlyR) onto pain fibers, reducing their ability to transmit pain signals to a higher center in the brain.

Chronic pain is a difficult-to-treat medical condition and new pharmacological targets are awaiting to be discovered. The most widely used analgesic class for pain includes NSAIDs (COX inhibitors, aspirin, ibuprofen, paracetamol, among many others). However, they are ineffective for chronic neuropathic pain and long-term use leads to gastric bleeding or perforation, and myocardial infractions.^3–8^ The second choice of analgesics are opioids, which are effective yet highly problematic due to their addictive properties and high mortality rate by overdose.^9–11^ Finally, gabapentin and pregabalin are widely used to treat neuropathic pain but growing concerns have been raised because of their misuse by drug abusers.^12,13^ All of these concerns collectively represent a therapeutic challenge that requires discovery of additional pharmacological targets and drug development efforts. Among the most recent targets to intervene in chronic pain, GlyT2 has emerged as a promising candidate.^3–6,14^ Therefore, insights into GlyT2 structure–function relationships and active screening of GlyT2 blockers are currently under development. Currently there are no FDA-approved drugs targeting the GlyT2 transporter. VVZ-149 is a dual GlyT2 and 5HT2A inhibitor, with a GlyT2 activity of 0.86 *µ*M, and is in clinical trials (2b stage), unfortunately, it shows low selectivity.^15^ Several other drug candidates did not advance to clinical trials because either they had a poor ADMET profile (ALX1393) or were irreversible binders (Org-25543), producing serious toxic effects by decreasing GlyT2 expression.^16^ Several drug candidates for schizophrenia, which inhibit GlyT1 homologs, failed for similar reasons.^17^ This has given researchers the impression that GlyT1/2 are not relevant targets. However, the correct balance of a number of factors may provide optimal drug candidates, which may replace current anti-chronic pain medicines. A good blood–brain barrier (BBB) ratio along with an optimal ADMET profile, reversible GlyT2 inhibition, and selectivity are the key factors for new modulators. For instance, a recently developed second-generation inhibitor GT-0198 showed excellent preclinical data.^18^ Unfortunately, this inhibitor exhibited also some, sometimes small, inhibition on other transporters.

The main problem during the design of all of the aforementioned inhibitors was the lack of high-quality structural and dynamics data, which can provide a deep understanding of the mechanism of action and selectivity. It has recently become evident that the B(0+) transporter (SLC6A14) is involved in the mTOR path and is greatly increased (up to 600) in pancreatic cell lines.^19^ The identity of the ligand binding domains (LBDs) between GlyT2 and B(0+) is 91% conserved; only three residues are different, from which presumably only one mutation, Ala284Ser, can modulate selectivity. Thus, inhibiting this transporter would greatly assist development of anti-cancer agents,^19^ but the side effects of inhibiting B(0+) during chronic pain treatment are unclear, and none of the known GlyT2 inhibitors have been tested for SLC6A14 activity. However, based on these data it is obvious that the selectivity against SLC6A14 cannot be neglected. On the other hand, structure-based design can provide lead structures for both pancreatic and chronic pain treatments, and such efforts continue in our lab.

For identification and characterization of new GlyT2 inhibitors in this study, we have undertaken an approach that combines state-of-the-art computational methodologies with biochemical assays to fully characterize the inhibitors. We paid special attention to the development of a high-quality GlyT2 structure that can be used for both hit identification and lead optimization, revealing ligands’ mechanism of action and selectivity. The identified new hit compounds should shed light on a novel family of molecules which, after proper optimization, can be tested in animal models of chronic pain.

## Results and discussion

### Generation of a high-quality GlyT2 structure by high-throughput molecular dynamics (MD) simulations

To obtain a high-quality GlyT2 structure that can be suitable for identification of new lead drug-like molecules by virtual screening (VS), we performed a series of intensive MD simulations. Based on the initial homology model (see SI Methods) we executed five independent 500-ns-long MD simulations on both the apo (without ligand) and holo (with ligand) conformations. Execution of multiple simulations ensures improved sampling and detection of the main structural binding pocket conformations. We obtained the center of the most populated cluster of structures (> 50% of the population) from all of the MD simulations by cluster analysis. Furthermore, this structure was energy-minimized for our *in silico* drug discovery study. We selected also an average structure for further examinations. Finally, 2×250-ns-long simulations on the serotonin transporter structure, pdb id: 5i6x, was performed (control runs). This allowed us to assess the quality of the MD studies with respect to the membrane parameters, *Na*^+^/*Cl^−^* coordination, and overall stability. Several MD simulations via a combination of Amber14SB and lipid 17 force fields were also executed and the structures were in close agreement to those obtained by CHARMM36 FF.

These simulations indicated that the GlyT2 transporter has a similar structure to those of serotonin, dopamine, and leucine transporters. We identified the amino acid residues that interact with glycine (Figures 1 and S1). The average root mean square deviation (RMSD) value of the backbone atoms between GlyT2 and the serotonin transporter was approximately 3Å, but this difference was mainly due to the rearrangement of the extracellular flexible loops. Residues interacting with glycine did not undergo any significant changes after the simulations were equilibrated (150 ns), which was confirmed by the good convergence observed (data not shown). Glycine, chlorine, and the major two *Na*^+^ ions were very stable along the simulations. The glycine carbonyl group forms H–bonds with the first sodium, Gly212, and Tyr287; whereas the nitrogen undergoes H–bonding with Ala208, Ser477, and Ser479. The *π*–*π* interactions with Tyr207, Phe476, and Trp482 are well-pronounced. The position of the third *Na*^+^ ion was revealed. It was coordinated mainly by Glu248, Met276, Trp263 and Glu648 and rapidly (after 50-ns of simulation) occupied this pocket (Figures S2). Our results are similar to those recently published by others^20^ and indicate that the *K*^+^ ion may also bind to the same pocket. It was coordinated by the same residues but with a position a bit shifted forward Glu248 (Figures S3A and S3B).

**Figure 1:**
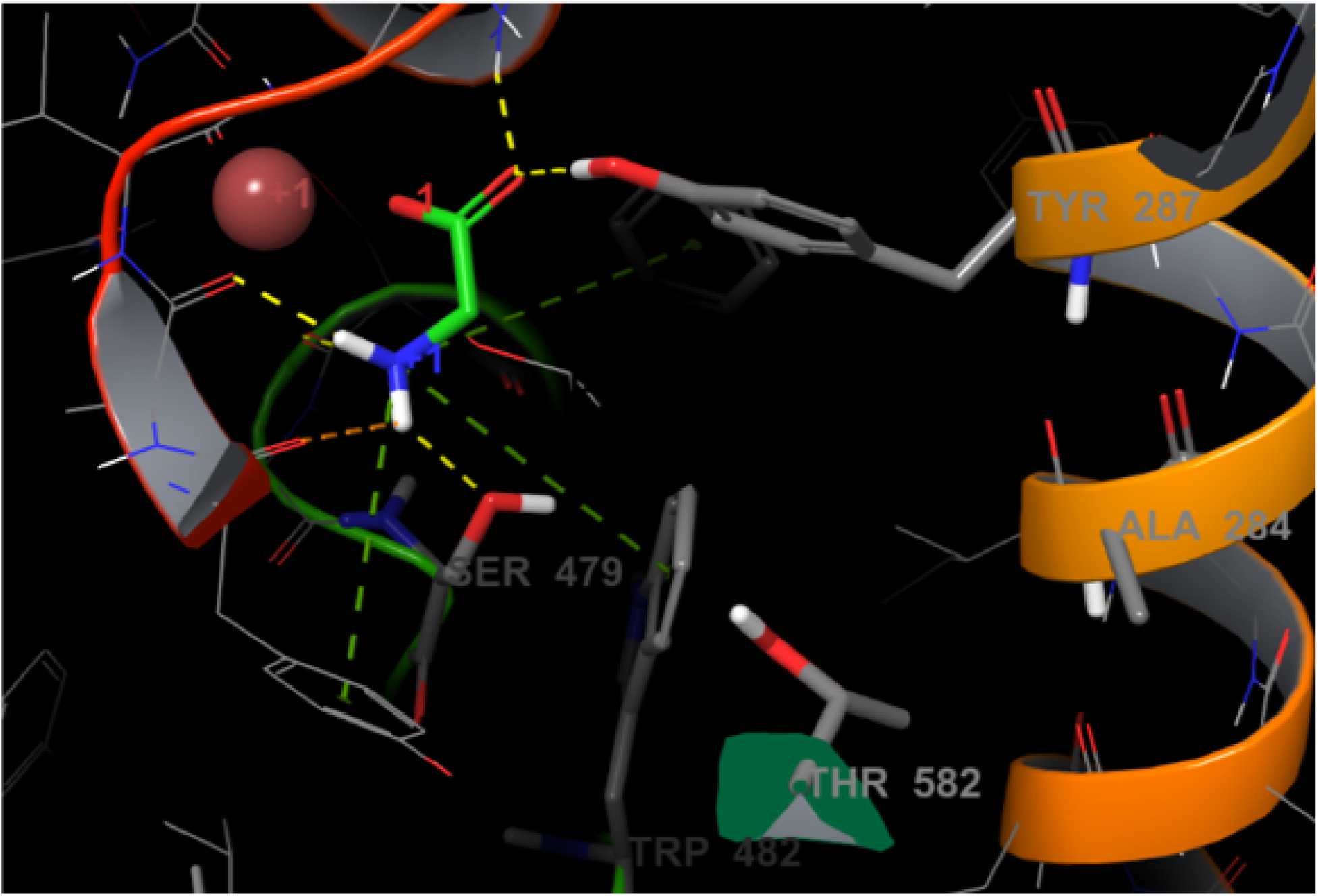
Glycine molecule in the binding site of GlyT2 as identified by our MD studies.

Our hypothesis is that the third *Na*^+^ ion is required for an additional glycine binding site stabilization, and in particular it affects the conformation of residue Trp482, critical for GlyT2 function and the residue’s neighboring amino acids. This is also supported by biochemical analyses which showed an increased glycine stabilization upon Glu248 and Glu648 mutations to hydrophobic residues.^20^ Stabilization via allosteric interactions of the second *Na*^+^ is also possible. To prove this we ran a 5x500-ns-long MD simulations on the GlyT1 transporter where the third *Na*^+^ is not present. The second sodium was less stable in GlyT1 and we even captured one of the likely unbinding paths in one of the MD runs (Figure S4). However, different paths are also possible.

We also studied the role of chlorine ion on stability. Destabilization of this ion via the Tyr233Phe mutation completely cancels the glycine activity. The lack of interactions between the chlorine and the OH group of Tyr233 led to a rapid dissociation of the ClËŮ ion which in turn destabilized the neighboring sodium and finally the glycine (Figure S5, Movie S1). At the end of these simulations both the first sodium and glycine left the transporter but the glycine moved out into extracellular space via its entry path, confirming that transport is impossible without chlorine.

### Impact of the individual residues to glycine binding as calculated by free energy perturbations (FEP+ protocol)

To quantify the residue contributions to glycine–GlyT2 interactions we conducted FEP+ calculations. We identified 10 residues which are important for compound binding (Table 1). Some of them are already known to significantly impact glycine activity. For instance, these include Ser477Ala, Phe476Ala, Ser479Ala and Trp482Ala, which completely abolish glycine transport.^21^ Our data match previously published qualitative and quantitative experimental studies. For instance, Ser479Ala decreased the EC50 value of glycine from 12 to 1070 *µ*M, a difference of G= 2.6kcal/mol; e.g., in a frame of the error of that predicted by the FEP+ value. This data indicates differences in the glycine binding affinity (G) upon mutations in the range of 1.5–4.0 kcal/mol, transforming glycine binding into the millimolar range; e.g., fully canceling it. Furthermore, the Tyr287Phe mutation weakens the glycine binding by 0.7 kcal/mol, due to the loss of one H–bond, which is in agreement with the commonly accepted strength of a typical hydrogen bond in a protein environment,^22^ suggesting that our FEP+ data and generated structure are precise. Thus, the preliminary FEP+ results provide a structural basis for the observed significance of the selected binding site residues to glycine transport. Most of these residues are conserved in GlyT1, dopamine, and serotonin transporters, yet some of them are unique to GlyT2, providing us helpful data on selectivity.

**Table 1:** Changes in the glycine binding activity (∆∆G; kcal/mol) upon 11 mutations as calculated by FEP+

| Mutation | Predicted $\Delta\Delta G$ | Predicted Error |
| --- | --- | --- |
| SER477ALA | 4.04 | 0.44 |
| SER479ALA | 3.37 | 0.43 |
| TRP482ALA | 2.62 | 0.45 |
| PHE476ALA | 1.58 | 0.43 |
| TYR207ALA | 1.43 | 0.46 |
| TYR287ALA | 0.97 | 0.44 |
| VAL209ALA | 0.89 | 0.41 |
| LEU211ALA | 0.69 | 0.42 |
| TYR287PHE | 0.69 | 0.42 |
| GLY212ALA | 0.44 | 0.79 |
| THR578ALA | -0.68 | 0.44 |

### Identification of new GlyT2 hit compounds as selective inhibitors by our novel VS protocol

We developed a special protocol for our high-throughput virtual screening (HTVS) with the aim of discovering new selective GlyT2 inhibitors. It has been established that a combination of the pharmacophore and docking methods provide superior lead identification success compared to employing a single approach.^23^ Currently, there are few known specific GlyT2 inhibitors. Their mechanism of action is diverse and these sets of ligands have low structural similarity. Thus, a ligand-based pharmacophore search was not suitable for our research needs. We used our newly developed in-house strategy that combines a recently refined structure-based pharmacophore approach and docking method in a frame of Schrodinger 2017-3 software (see SI Methods) [ref 4 in SI]. This type of pharmacophore model is based on the docking of several hundred fragments into the binding site and the pharmacophore points are defined in accordance with their best position and binding score. These are the places within the binding site that were predicted to be essential for ligand binding.

We generated four pharmacophore points, requested four of four matching points, and screened over 3.5 million compounds from the ZINC15 library (Figure 3).^24^ Point D17 (donor) represents the requested H–bond with Ser479, which is, as it was shown above, important for selectivity and is mutated to Gly in GlyT1. Furthermore, we subjected 35,000 best candidates to docking and visually inspected the top-scoring 100 ligands. The top 20 ligands were selected by visual inspection and re-docked then by induced-fit docking (IFD). The IDF procedure was accomplished by metadynamics refining the binding modes as shown previously (Figure S6) [ref 6 in SI]. Two different structurally similar groups of ligands have been found to be most promising GlyT2 inhibitors (Figures 2A, 2B). We selected four of them (two from each group) for biological testing (Figure 4).

**Figure 2:**
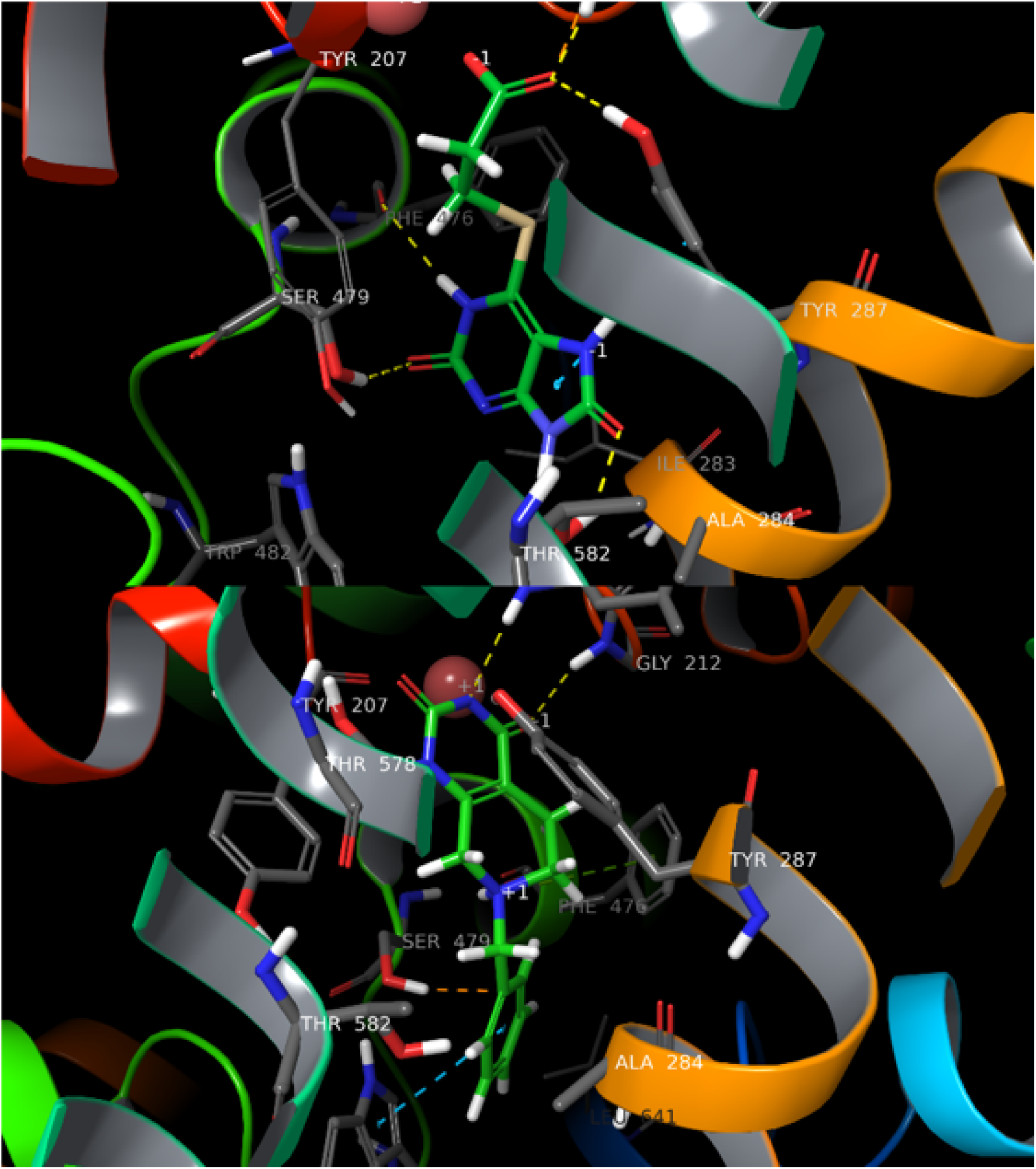
A) Lead 1 and (B) Lead 2 in the binding site of GlyT2, identified by HTVS and MD studies.

**Figure 3:**
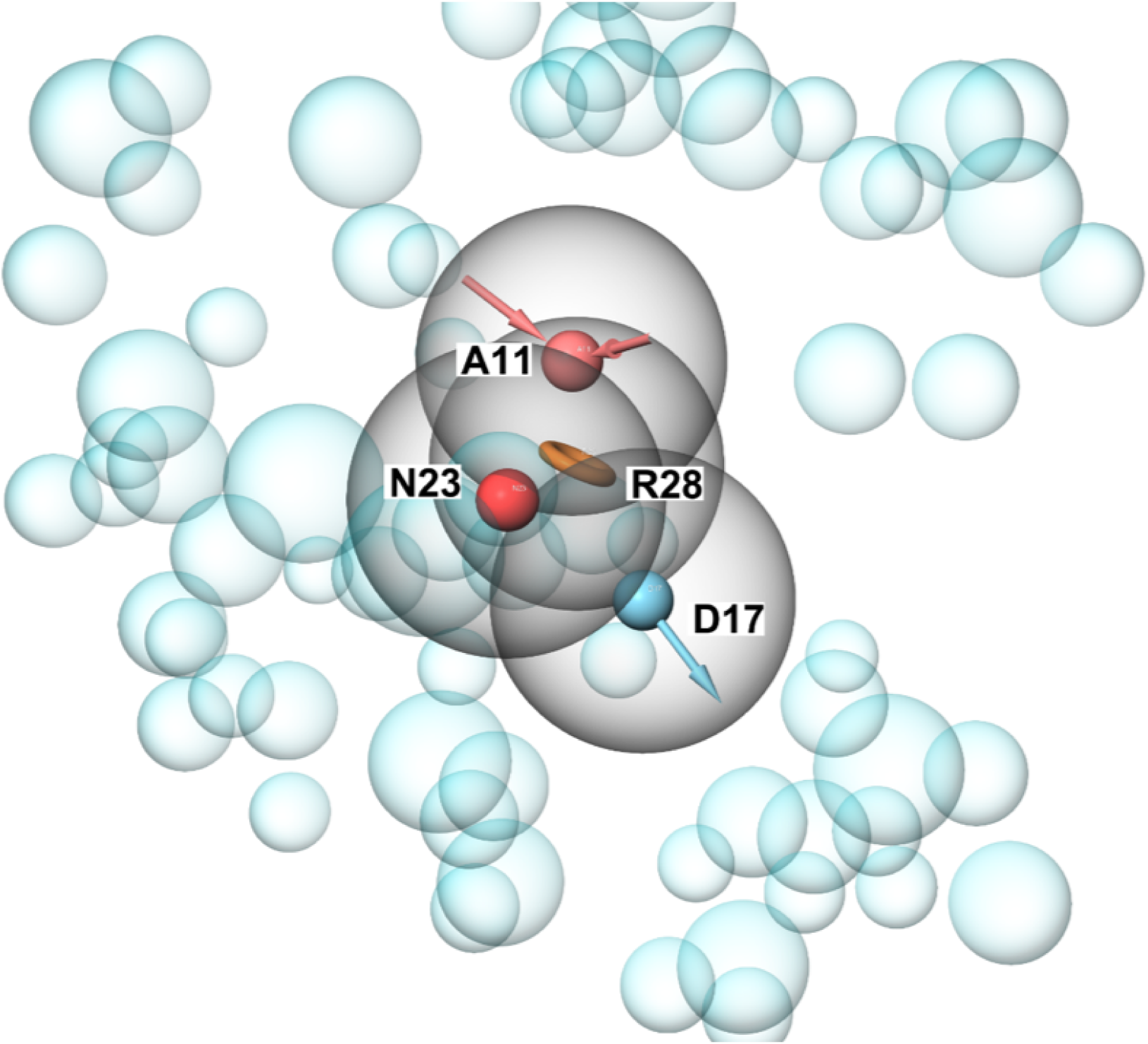
Structural based pharmacophore model based on the GlyT2 MD generated structure. Four pharmacophores (Acceptor (A11), Negative charge (N23), Aromatic ring (R28) and Donor (D17)) were used.

**Figure 4:**
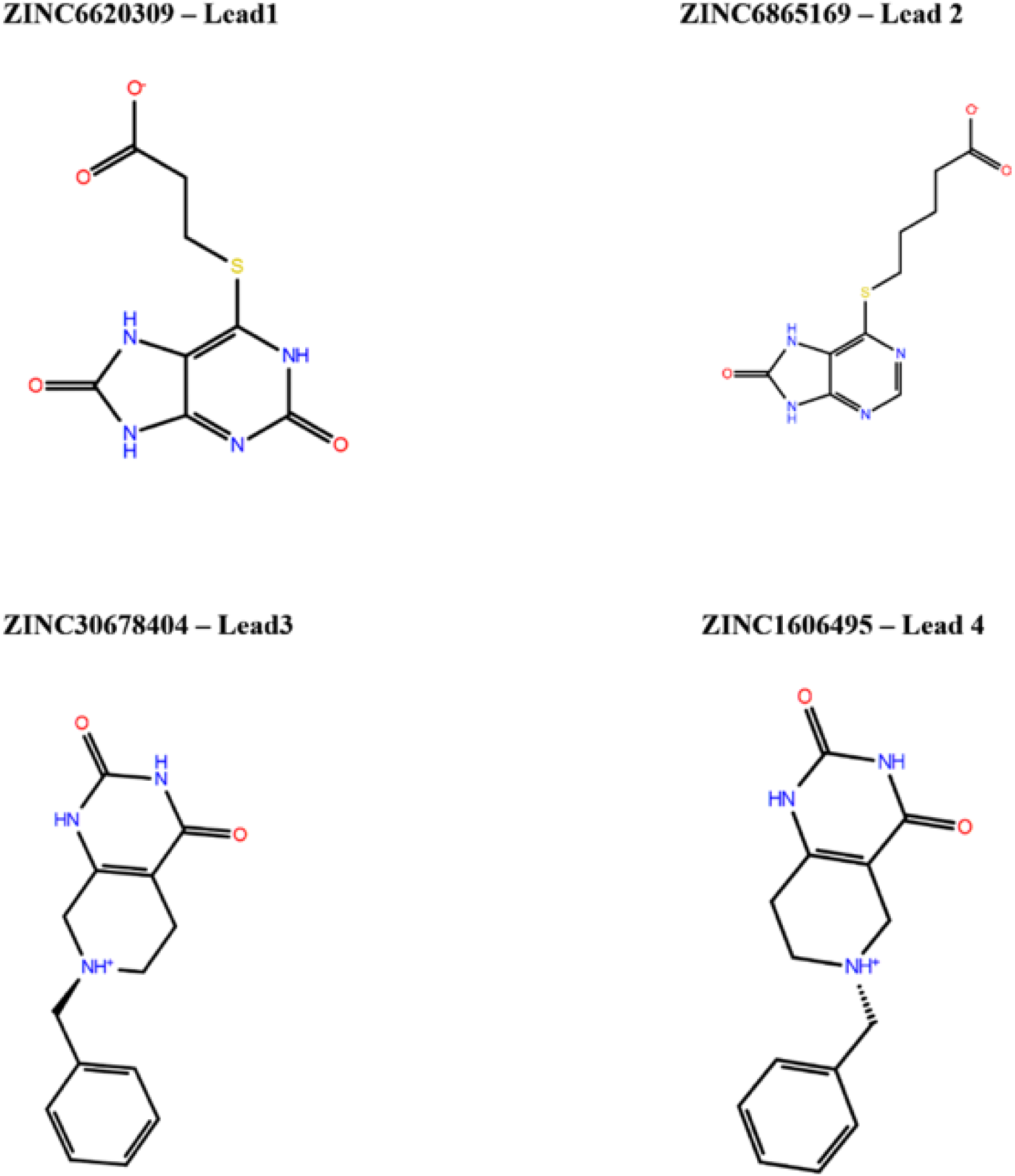
Identified by VS hits which were confirmed by in vitro studies.

### Inhibition of GlyT2 activity by compounds obtained by *in silico* screening

To validate the use of our in vitro assay for screening purposes, we initially carried out inhibition studies with the selective GlyT1 inhibitor ALX5407 and GlyT2 inhibitor ALX1393.^25–27^ The inhibitor ALX1393 has been the most extensively studied in different models of pain in rodents, shows 40-fold higher selectivity for GlyT2 over GlyT1, and inhibits the human GlyT2 at nanomolar concentrations. To investigate the response of the selected compounds, we used porcine aorta epithelial cells stable expressing the human GlyT2. In these cells, the transporter traffics and accumulate at the plasma membrane where shows a Km of 123 *µ*M and Vmax of 8.2 nmol/min/mg (Figure S7). In control experiments shown in Figure 5A, we measured the concentration dependence of ALX1393 and observed a rapid inhibition of transport, with an average IC_50_ = 25.9 nM, a value which is in an agreement with literature data.^26,27^ This observation confirms that our model system expresses a functional human transporter which transports glycine with the expected specificity, and the activity can be inhibited by the traditional inhibitor ALX1393. As expected, incubation with the GlyT1 inhibitor ALX5407 showed a slow rate of inhibition with a calculated IC_50_ = 1.8 *µ*M (data not shown).

**Figure 5:**
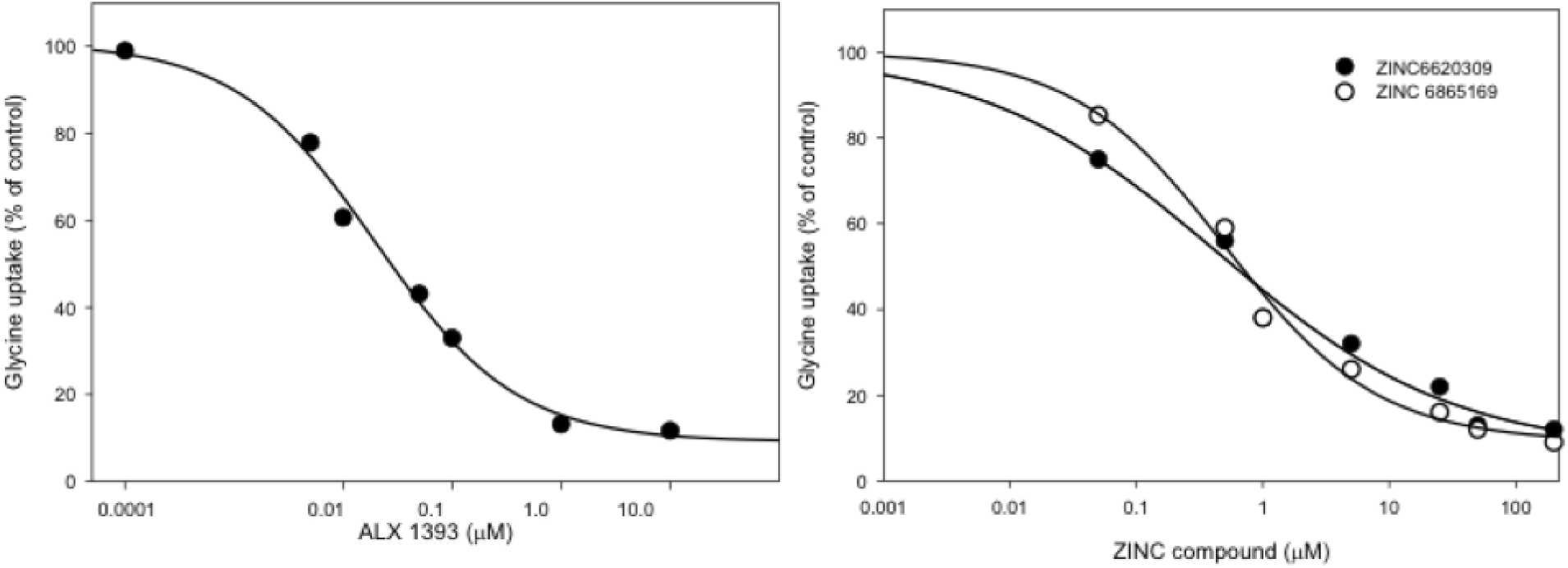
Inhibition curves of GlyT2 activity by ALX1393, ZINC6620309, and ZINC 6865169. We incubated PAE cells expressing GlyT2 with varying concentrations of inhibitors in the uptake mix and measured the activity for 10 min. We obtained sigmoidal curves and calculated IC_50_ values via Sigma Plot.

Furthermore, we tested four selected compounds identified by *in silico* screening (Figure 4). Interestingly, a small lead molecule 1, termed Lead1 (Figures 2A, 4 and 5; compound 1, ZINC6620309), consistently inhibited the transport at high nanomolar concentrations and showed an IC_50_ = 0.48 *µ*M. This compound was also greatly selective over GlyT1 (IC_50_ > 200 *µ*M). The structural analog, Lead2, which consists of two more carbon atoms after the propionic acid segment, and modification of the aromatic ring, was also active but featured slightly decreased (0.52 *µ*M) inhibition; i.e., still good inhibitor. This is presumably due to the longer chain that cannot be accommodated well in the ligand binding domain (LBD) and/or lack of significance for the binding interactions with Ser479 (Table 2). The second class of ligands was also active. Lead 3 showed an IC_50_ = 33.2 *µ*M. This is because that simple compound provides only interactions close to the first *Na*^+^ binding pocket where the GlyT1 and 2 are in practice identical (Figure 2B). Finally, Lead 4 was not potent due to the position of its long chain, providing less access to the key GlyT2 residues. In conclusion, these data collectively demonstrate that our structure is of high quality, allowing us to discover a new set of compounds with good inhibitory properties that can be further optimized. Based on the aforementioned data, we chose Lead1 as our primary candidate for further lead optimization and studied this ligand in more details.

**Table 2:**
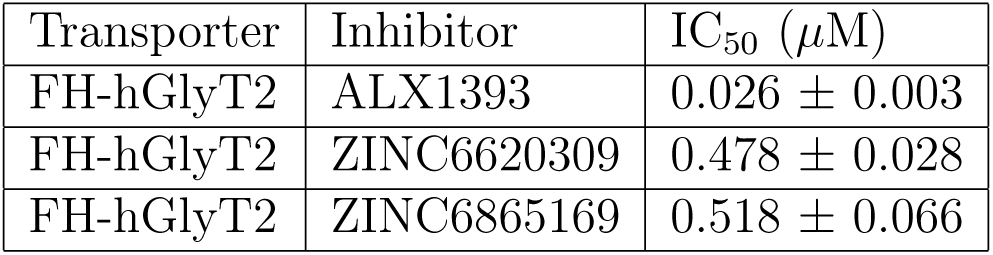
Inhibition of glycine uptake by ZINC compounds in cells expressing the human GlyT2. The IC_50_ value was determined by varying the concentration of inhibitor from 0 to 10 or 100 *µ*M at pH 7.5. Values are the average of three to five determinations with an average standard error of less than 12%.

### Structural basis of the observed Lead1 and 2 activities by MD

We procured relaxed Lead1–GlyT2 and Lead2–GlyT2 complex structures by executing 2x250 ns MD simulations via Amber 16. The carbonyl group of Lead1 interacts with the first sodium, Leu211, Gly212, and Tyr287. Based on this structure it was evident that Tyr287, Ser479, and the backbone oxygen of Thr578 makes stable H–bonding with the ligand and stabilize its aromatic component, in a conformation which provides *π*–*π* stacking with Phe476 and Trp482 (Figure 2A). We also observed a stable H–bond between Lead1 and the Phe476 backbone oxygen. Thr578 coordinated the second sodium but also can flip its -OH group to form an H–bond with Tyr287, thus accelerating the *Na*^+^ unbinding dissociating and demonstrating that Lead1, like ALX1393, is presumably a reversible inhibitor. Both Ser479 and Thr582 are dynamic in the presence of Lead1. Thr582 can form H–bonds with the ligand, the backbone of Thr578, and Tyr207, whereas Ser479 can do so with Lead1 and Trp482. These variations are due to the significant solvation of the LBD which was formed by the ligand’s multiple polar groups. Furthermore, we confirmed these results during our FEP+ guided lead optimization by execution of different sets of Lead1 transformations (unpublished data). The percentage of H–bond formation of Lead1 with Thr582 and Ser479 varied between 25% and 65% for both of these residues (Figures 6A and 6B).

**Figure 6:**
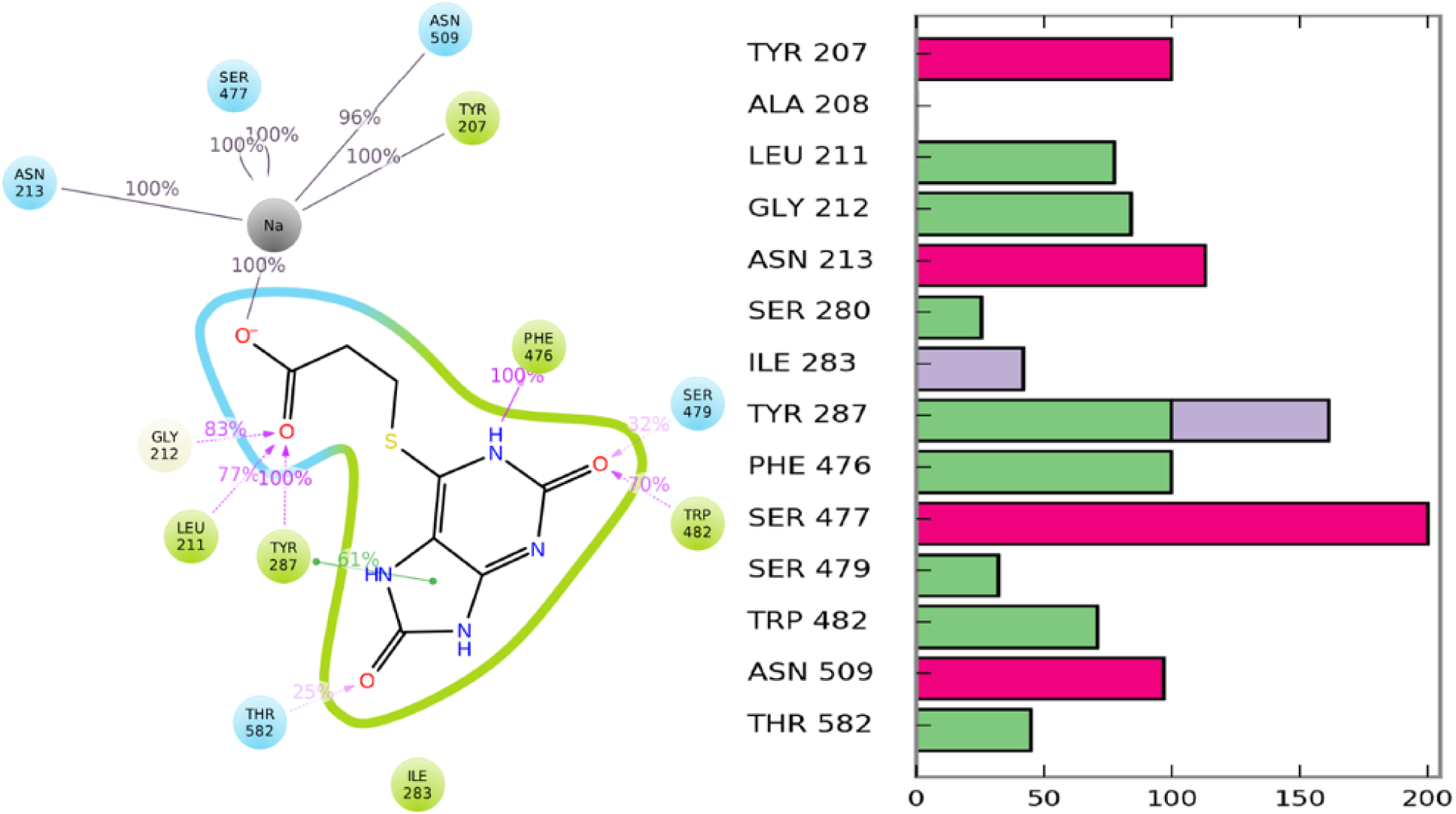
Identified Lead1 binding mode and Lead1–GlyT2 interactions, during the FEP+ calculations, represented by (A) a 2D view and (B) presence of GlyT2 interactions with GlyT2 residues. Purple and green lines show H–bonds and *π*–*π* stacking, respectively; whereas green, red, and blue bar charts indicate H–bond, ionic, and hydrophobic interactions, respectively.

Similarly, the Lead2 pyrimidine group interacts with residues close to the first sodium but provides limited connection with the *Na*^+^ ion itself. The phenyl ring facilitates *π*–*π* stacking with Phe476 and Trp482, but only hydrophobic interactions with Ser479 and Thr582 were observed (Figure 2B). Instead, these interactions were more pronounced with Ile283, a residue which is conserved between GlyT2 and GlyT1. We obtained similar results for compound 4, yet we observed one additional H–bond with Phe476 and stacking with Tyr287 (Figure S8). These results provide structural basis of the observed lower Lead2 selectivity.

### Lead1 selectivity as calculated by FEP+ and ADMET profile

Despite the high sequence similarity between the GlyT2 and GlyT1 binding sites, the experimental data for our discovered Lead1 compound indicated that it is a highly selective GlyT2 inhibitor (IC_50_ of 0.48 *µ*M) and the GlyT1 activity was >200 *µ*M. The difference between the binding sites of these subclasses of transporters is only five residues. Thus, Lead 1 is an ideal compound for revealing the selectivity. In GlyT1, the Ser479 and Thr582 residues are mutated to Gly and Leu, respectively, whereas Phe476 is mutated to tyrosine. Thus, it is clear that these mutations would have a great impact on Lead1 selectivity.

To quantify the contribution of all five mutated residues in GlyT1, and in particular Ser479Gly and Thr582Leu, we executed FEP+ ligand selectivity calculations. We employed our recently improved FEP+ sampling protocol^28^ which has been shown to significantly improve the FEP+ results. Moreover, an additional equilibration in the pre-REST stage of FEP+ on that system should be performed due to the structural rearrangements introduced by the mutations in the aforementioned GlyT2 structure. Indeed, the Ser479Gly mutation negatively impacted the selectivity and contributed G=+0.98 kcal/mol, whereas the Thr582Leu had the most significant contribution, G=+2.29 kcal/mol (Table 3). This is in an excellent agreement with the recent mutations studies for both ALX1393 and glycine.^29^

**Table 3:**
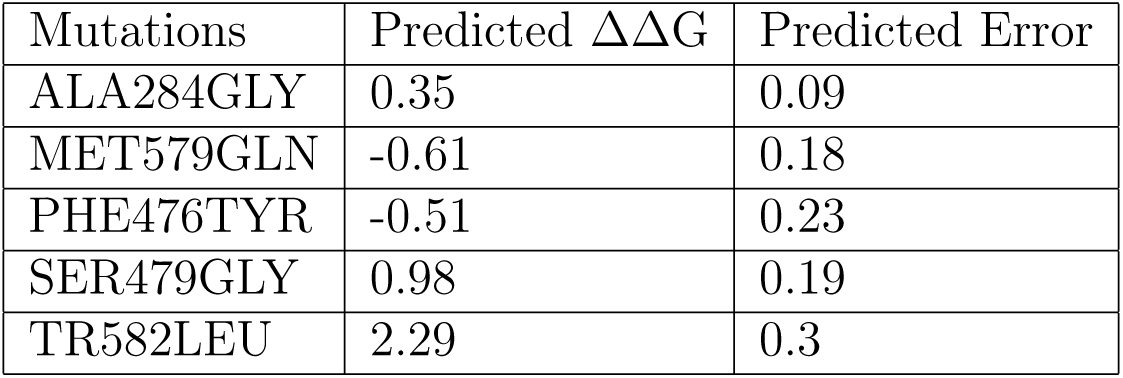
Changes in the Lead1 binding affinity (∆∆G; kcal/mol) upon the five mutations constituting GlyT2/1 differences as calculated by FEP+

It has been shown that the Ser479 to Gly mutation decreased glycine affinity form 13 to 72 *µ*M, which corresponds to 0.95 kcal/mol (G=0.59lnKd); e.g., the same value calculated as per our FEP+ simulations. Further, a significant decrease in the inhibition for the same mutation was also observed in ALX1393, whereas Thr582Leu completely abolished the glycine transport.^29^ This is an indication that in the glycine binding site all of these ligands facilitate similar interactions, which is reasonable result considering their similar structure at this part of the GlyT2 LBD: a carbonyl group and donor placed close to the 1st sodium and Ser479, respectively. However, for the remainder of the LBD it seems that ALX1393 and Lead1 exhibit different interactions. Additional support comes from the data for Lead 2 (Figure S9). Essentially, it provides ligand–protein interactions similar to those of Lead1 but the contact with Ser479 was not so significant. As a result, the selectivity was mainly due to the Thr582 H–bond. Notably, our FEP+ calculations showed that the Ala284Gly mutation had a small negative effect for the Lead1 activity whereas Phe476Tyr had a positive effect. In contrast, the last mutation abolishes the Glycine transport and reduces ALX1393 binding yet does not cancel it.^29^ Thus, Lead1 interactions in this part of the LBD are different than those of ALX1393.

Interestingly, in the B(0+) (SLC6A14) transporter Ser479 and Thr582 are unchanged, and Phe476 is changed to tyrosine. Assuming that the Ala284 is mutated to serine in the same transporter, which is in a close proximity to the Lead1 oxygen (see Figure 2A), we propose that Lead1 would be an even stronger inhibitor of B(0+). Thus, additional lead optimization can lead to discovery of both chronic pain and pancreatic agents.

In summary, these results provide a structural basis to the experimentally observed Lead1 selectivity in GlyT2, and additional support to the idea that this ligand cannot be expected to have more than sub-millimolar activity in GlyT1. An important outcome from the FEP+ calculations was also an understanding of how more-selective GlyT1 ligands can be discovered. For instance, it is evident that the Thr582Leu substitution would greatly increase the hydrophobic nature of the GlyT1 cavity, whereas the Ser479Gly substitution will cancel the acceptor possibility in this segment of the binding pocket.

The ADMET profiles of the compounds were computationally predicted using QikProp (see the attached excel sheet in Supporting Information). Lead 1 shows poor membrane permeability by our calculations. We also speculate that due to the presence of the carbonyl group in Lead1 as a key functional group in the interactions with the first sodium ion and the binding as a whole, as in the case of ALX1393, this inhibitor will be reversible and thus will not produce undesired toxic affects due to decreasing GlyT2 expression levels.^16^ However, the calculated Lead2 ADMET profile was superior, indicating superior BBB penetration and no significant interactions with major proteins that can produce side effects. However, additional selectivity screening against the remaining members of these transporters and an additional optimization will be necessary. We are continuing these efforts toward improved drug development for chronic pain management.

## Supporting information

see the excel sheet

Moive S1

## Acknowledgement

We thank the High-Performance Computing support staff (Marc T. Hertlein and Leopoldo Hernandez) at The University of Texas at El Paso for assistance in using the Chanti cluster. We acknowledge the Texas Advanced Computing Center (TACC) at The University of Texas at Austin for providing HPC resources that have contributed to the research results reported within this paper. URL: http://www.tacc.utexas.edu. We also thank Ricardo Avila and Michael Scott Long for reading and editing the paper.

## Author Contributions

SS, FF and MM conceived the project. SS supervised the computational part of the project and MM supervised the biology part of the project. FF performed all the Computational Work. EP Performed the biological work. FF, SS, and MM contributed to the analysis of results and wrote the manuscript. All authors reviewed and edited the manuscript.

## Supporting Information Available

The following files are available free of charge.

- SI_Methods.pdf: Contains methods, additional figures and references
- qikProp_ADMET_predictions.xlsx: computational ADME predictions of 4 listed compounds
- GlyT2_Movie_S1.avi: MD simulation movie of glycine unbinding mechanism in GlyT2.

