## Supplementary material for "Discovery of New Classes of Glycine transporter 2 (GlyT2) Inhibitors and Study of GlyT2 Selectivity by Combination of Novel Structural Based Virtual Screening Approach and Free Energy Perturbation (FEP+) Calculations": SI_Methods.pdf

**2 Figures** **7**

**References** **18**

### 1 Methods

#### 1.1 Homology modeling

A detail analysis of all possible homologous structures to GlyT2 was initially performed. For our homology modeling we used both the Prime package of Schrodinger software and I-TASSER server. Recently obtained high resolution X-ray structures of *Drosophila* dopamine (pdb id: 4xpt) and human serotonin (pdb id: 5i6x) transporters were employed as templates [1, 2]. They shared higher similarity compared to the older bacterial (*Aquifex aeolicus*) structure of leucine receptor (pdb id: 3f3a). The later showed only 22% identity to GlyT2 whereas the pdb id 4xpt and 5i6x structures have 42% and 38% similarity, respectively. For the aim of our study, the identity of amino acids in the glycine binding site is also very important. For instance, the serotonin transporter has a 68% identity with GlyT2, which was much better than that of the bacterial leucine transporter (41%). In summary, several of the previous models of glycine transporters have been based on the leucine receptor structure but they should be used with caution. For instance, pdb id 3f3a is populated with more than one ligand, which can produce significant changes in LBD, and also the similarity of helix 12 (H12) is less pronounced to GlyT2 than those of serotonin and dopamine transporters.

#### 1.2 Virtual screening approach

The structural and ligand-based *in silico* screens are of great help during drug discovery nowadays, and their combination has been shown to provide better results than using a single approach. However, many targets have not had any ligands discovered, thus making this combination impossible.

In the current study we tested whether the recently introduced structural based pharmacophore approach in Phase software (Schrödinger package) in combination with docking followed by rescoring with the new Glide-SP scoring function can achieve reasonable virtual screening (VS) results. These methods were also tested in our lab on the full (102 targets) DUD-E benchmark. Surprisingly, we found that the structure based pharmacophore hypothesis created by the docking of more than 600 small fragments (the default number in Phase) performs well for most of the targets with an average active ligand recovery rate of 10-15% during the first 10% database screen [3] (see Figures S10 and S11 for examples). The MD prepared structures further increased these values. These results are equal to the ligand based pharmacophore virtual screening. The docking technique performed much better on DUD-E set. Early enrichment performance shows on average of about 30% of known actives are recovered in screening the top-ranked 1% of recovered decoys, similarly to the previous DUD-E test results. Further, an improvement of the Glide-SP scoring function, as it is also claimed on Schrödinger website, was confirmed as it is shown on the ACE test target, which usually produced unsatisfied results previously (Figure S12). However, the main aim of a real drug discovery project is the very early enrichment performance, i.e. how many active compounds will be discovered between the first 10 to 50 ligands, which we call here also success rate (SR) or efficiency (EF). According to our data the combination of these methods achieves an impressive result of nearly 75% success rate for all 102 targets, even reaching often 100%, for the top 10 ligands [3]. Based on our data we developed a new VS protocol using the top 10% of the pharmacophore search (3.5 million ZINC compounds in our case) as an input to the docking and scoring by Glide-SP (35,000 ligands). All calculations were performed by the Schrodinger's Phase and Glide software packages as implemented in Schrodinger 2017-3 suite [4]. We hope that our approach will be in help during the drug discovery projects and in particular in a case that no know ligands are available for the studied protein. The aforementioned short description provides only some brief introduction to our approach, and a detailed paper about the topic is underway. However, as this methodology and results has been based on X-ray data [3], for us it was also interesting to find out to which extent it would be applicable for generated by homology modelling structures as in the case of GlyT2 transporter. For more description see also the results and discussion part of the paper.

#### 1.3 Pharmacophore, regular and induced fit (IFD) docking

As it was described above we used the combination of structure-based pharmacophore and docking approaches as they are implemented in Schrodinger Suite 2017-3. For both of these methods the default settings were employed. Exceptions were that during pharmacophore search all points match (4 from 4 pharmacophores (Figure 4)) were requested whereas during the docking we kept 10 000, instead of 5000, initial docking poses and the rewards of H-Bonds were employed as an option. The induced Fit Docking (IFD) [5] were also performed using the default settings but either an increased accuracy to XP docking mode or enhanced sampling option were also used to test results stability. This was done in order to investigate if other docking poses can be obtained when using a different settings.

#### 1.4 Metadynamics refinements of IFD solutions

Metadynamics refinements of the IFD output possess is an important step in order the correct input pose to be found. Very often the regular IFD suggests variety of docking solutions which are similar to two major conformation of the ligand; in our case the aromatic ring conformation rotated by 180 degree. On the other hands the advanced MD sampling techniques, such as accelerated MD (aMD) and metadynamics, provide a good opportunity the most likely binding mode to be recovered [6, 7]. In current study we used recently suggested approach based on execution of several independent metadynamics simulations and then scoring the ligand docking poses by observed ligand-protein interactions during the simulations course [6]. This protocol is now implemented in Schrodinger suite and can be automatically performed. We used the default settings for these calculations. Essentially, 8 independent 10-ns-long metadynamics simulations were executed for selected two possible conformations and the pose scores were retrieved as previously suggested (Figure S6) [6].

#### 1.5 Molecular dynamics

All molecular dynamics (MD) simulations were carried out using the Amber 16 program [8]. CHARMM36 force field (FF) was used for the calculations of the apo transporter form and those bound with glycine (holo form). The GlyT2 was enabled embedded in a POPC membrane by the CHARMM-GUI membrane builder webserver (charmm-gui.org) [9]. Membrane orientation was obtained by OPM server [10] and was essentially same as for serotonin, which was used also for a control run, and dopamine transporters. Default parameters were used for both heterogeneous lipids creation. Water thickness of 17.5Å and 0.15M NaCl ion concentration were used. Simulations with NaK were also provided in order to examine potential difference in $Na^+$ and $K^+$ at the $3^{rd}$ sodium binding site. The system size was X=99 Å, Y=99 Å and Z=152 Å and it contained about 98 000 atoms for both the apo and holo forms. The glycine parameters as calculated by CHARMM General Force Field (CGenFF) were employed. We used the default relation times, as generated for Amber software, by CHARMM-GUI but used friction coefficient of 2 ps-1 in order to allow higher temperature (T=303.03K) fluctuations. The non-bonded cutoff was set to 12 Å whereas the force-based switching was 10 Å. For these calculations the MC Barostat was used for the pressure control. After 8000 steps minimization (2500 with the conjugate gradient method), five equilibration steps were executed and five independent 500-ns-long production runs were performed for each system. Finally, based on all of these 2.5$\mu$s MDs we retrieved both average and cluster (center of the most populated cluster) structure for both apo and GlyT2 transporters, which were used for our further analysis, VS and FEP+ calculations. However, for identified lead compounds we executed simulations with Amber14SB, along with Lipid 14, and GAFF2 force fields [8]. The reasons were: 1) comparison of the structures produced by different FFs, which would provide more confidence about the quality of obtained structures and 2) our experience shows that GAFF2 FF handle more complicated compounds better. Another difference is that for these calculations we used Berendsen instead of MC Barostat because of our recent results [7] showing that the MC Barostat should be used with caution for ligand sampling in similar systems. For these MD runs, the systems were energy-minimized in two steps. First, only the water molecules and ions were minimized in 6000 steps while keeping the protein and ligand structures restricted by weak harmonic constrains of 2 kcal mol–1 Å–2. Second, a 6000 steps minimization with the conjugate gradient method on the whole system was performed. Furthermore, the simulated systems were gradually heated from 0 to 310 K for 50 ps (NVT ensemble) and equilibrated for 3 ns (NPT ensemble). The production runs were performed at 310 K in a NPT ensemble. Temperature regulation was done by using a Langevin thermostat with a collision frequency of 2 ps–1, and the pressure regulation via Berendsen barostat. The time step of the simulations was 2 fs with a non-bonded cutoff of 8 Å using the SHAKE algorithm [11] and the particle-mesh Ewald method [12]. Two independent 250-ns-long production simulations were executed for all of the lead compounds complexed with GlyT2.

#### 1.6 Free energy perturbation FEP+ calculations

All calculations have been conducted using the Schrödinger molecular modelling suite 2017-3 [4]. Free energy perturbation calculations were performed using the FEP+ methodology, which combines the accurate modern OPLS3 force field [13], GPU-enabled high-speed molecular dynamics simulations with Desmond MD package [14] , the REST algorithm for locally enhanced sampling [15], a cycle-closure correction [16] to incorporate redundant information into free energy estimates. The FEP+ calculations were based on the aforementioned MD obtained structures and were conducted using the default protocols: The systems were solvated in an orthogonal box of SPC water molecules with buffer width (minimum distance between box edge and any solute atom) of 5 Å for the complex and 10 Å for the solvent simulations. The full systems were relaxed and equilibrated using the default Desmond relaxation protocol, consisting of an energy-minimization with restraints on the solute, then 12 ps length simulations at 10 K using an NVT ensemble followed by an NPT ensemble. A total of 12 $\lambda$ windows were used for all calculations. Replica exchanges between neighbouring $\lambda$ windows were attempted every 1.2 ps. Finally, for the actual FEP+ pre-REST and REST calculations we employed our new FEP+ sampling protocol which demonstrated superior results especially with MD derived structures [7, 17] . Thus, for these FEP+ simulations we employed 2x10 ns/$\lambda$ pre-REST and 8 ns/$\lambda$ REST calculations. The convergence was closely monitored. All calculations were run on Nvidia Pascal architecture GPUs (cluster of 8xGTX 1080Ti GPUs).

#### 1.7 ADMET

All ADMET calculations were performed by QikProp as implemented in Schrödinger suite 2017-3 [4]. QikProp predicts the widest variety of pharmaceutically relevant properties - octanol/water and water/gas log Ps, log S, log BB, overall CNS activity, Caco-2 and MDCK cell permeabilities, log Khsa for human serum albumin binding, and log $IC_{50}$ for HERG $K^+$-channel blockage - so that decisions about a molecule's suitability can be made based on a thorough analysis. QikProp bases its predictions on the full 3D molecular structure; unlike fragment-based approaches, it can provide equally accurate results in predicting properties for molecules with novel scaffolds as for analogs of well-known drugs.

#### 1.8 Development of an in-vitro cell system for screening of small molecules

To set up a cell assay for screening and testing of GlyT2 inhibitors, we cloned the human GlyT2 gene into the expression vector pcDNA3.1 and used it for transfection and selection of porcine endothelial aortic (PAE) stable cell lines. We selected PAE cells as our preferred model cells because the cells firmly attach to the plastic and do not overexpress ectopic proteins, Moreover, they are flattened which allow easy visualization of the plasma membrane by conventional fluorescence microscopy. After selection of stable cell lines, we analyzed the localization of the transporter by immunofluorescence with anti-GlyT2 specific antibodies. As shown in Figure S7, GlyT2 expression in PAE cells shows localization of transporter at the plasma membrane, particularly concentrated in ruffle- and phylopodia-like structures. Staining of the cells with DAPI showed that the majority of the cells express GlyT2. By contrast, immunostaining of GlyT2 expressing cells with anti-GlyT1 antibody, a homolog transporter, showed no staining around DAPI-containing cells demonstrating an axenic and homogenous cell line. In addition, we have established cell line expressing the human GlyT1b that has been fully characterized and shown to express a functional GlyT1b [17, 18]. As demonstrated in the published manuscripts and shown in Figure S9, GlyT1b cells stained with antiGlyT1 antibody identified transporter mostly at the plasma membrane but no reactivity was obtained with the anti-GlyT2 antibody. To confirm the presence of functional transporters in PAE cells, we measured [3H] Glycine uptake over different concentrations of substrate (Figure 4B). The cell line exhibited specific transport defined as the difference between [3H] Glycine uptake from GlyT2 cells minus parental PAE. This kinetic analysis shows a Michaelis-Menten behavior with an affinity for glycine of 12.3 $\mu$M and a Vmax of 8.2 nmol/min/mg (Figure S7). These results together validate that this cell line has the correct localization and good quantities of functional GlyT2, making this cell line ideal for screening of a novel class of specific inhibitors.

#### 2 Figures

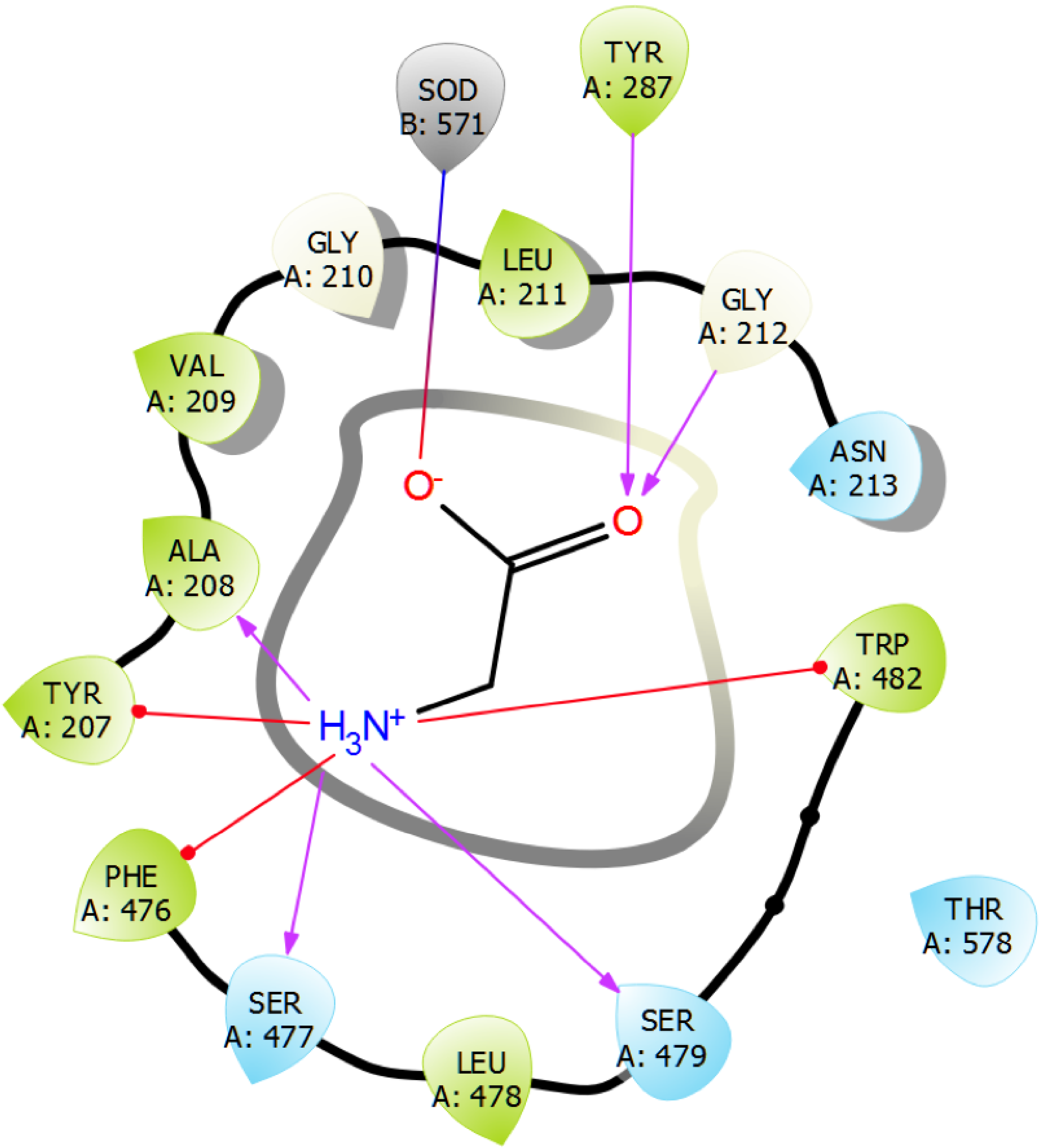

Figure S1: 2D representation of Glycine-GlyT2 interactions as identified by our MD studies.

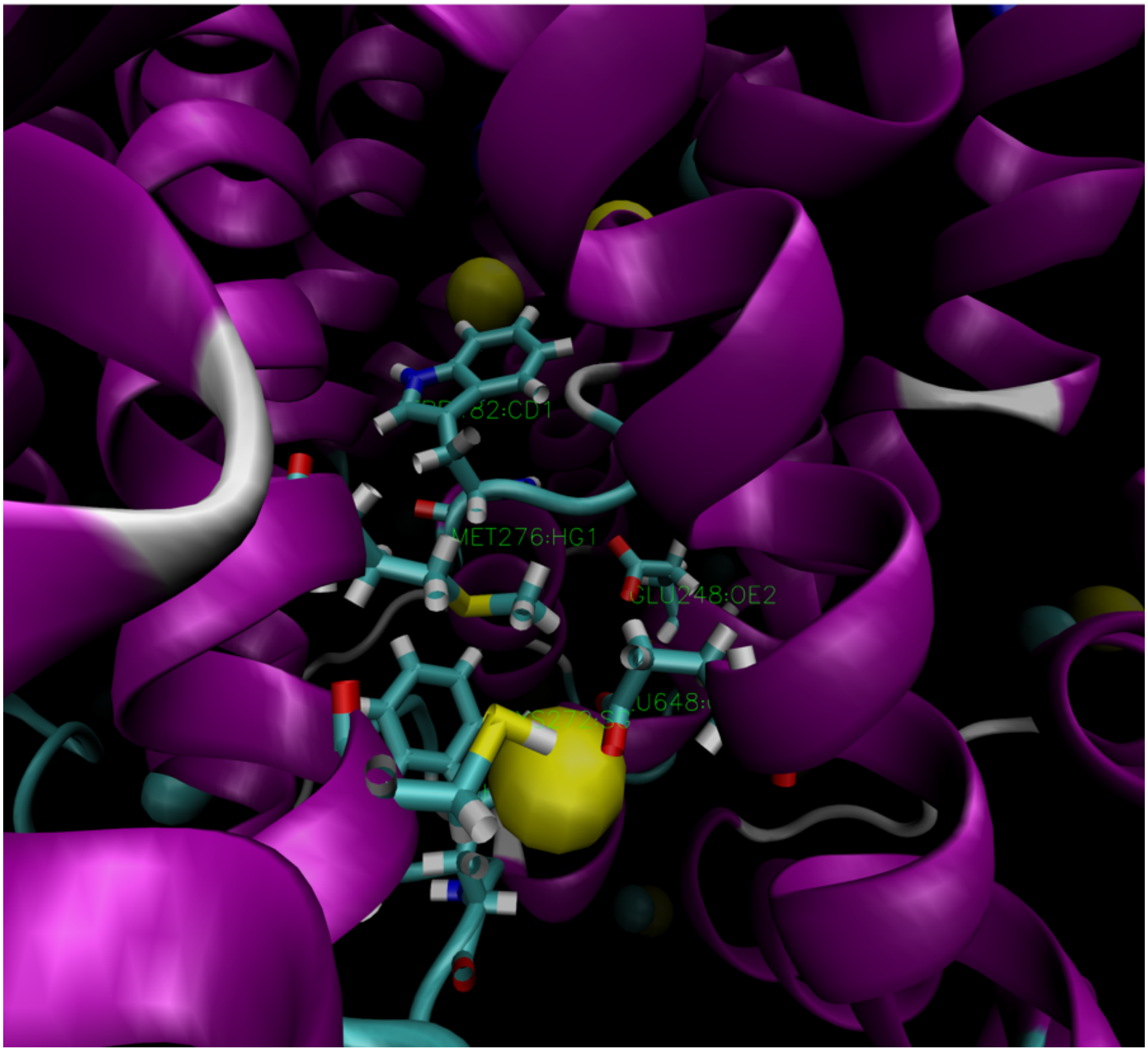

Figure S2: 3D representation of the $3^{rd}$ sodium binding site.

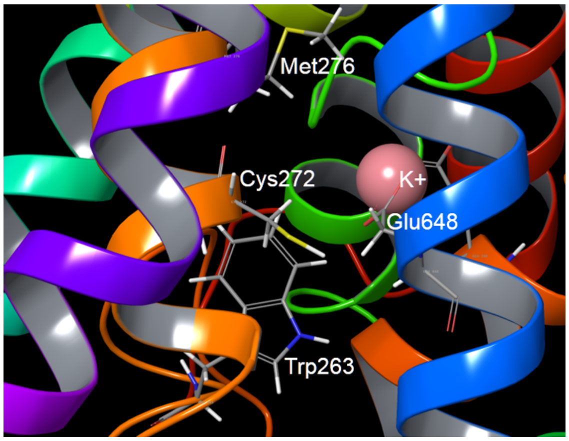

(a) 3D represntation

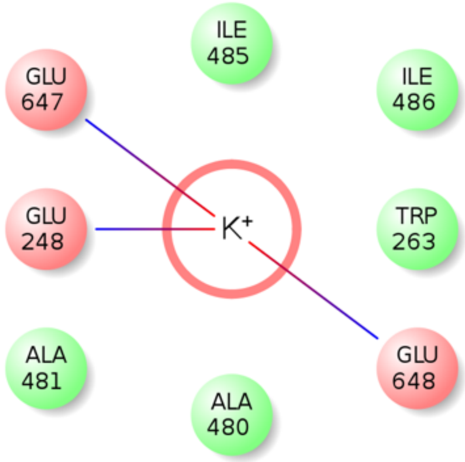

(b) 2D representation

Figure S3: (a) 3D and (b) 2D representation of the $K^+$ ion position in the $3^{rd}$ sodium binding site.

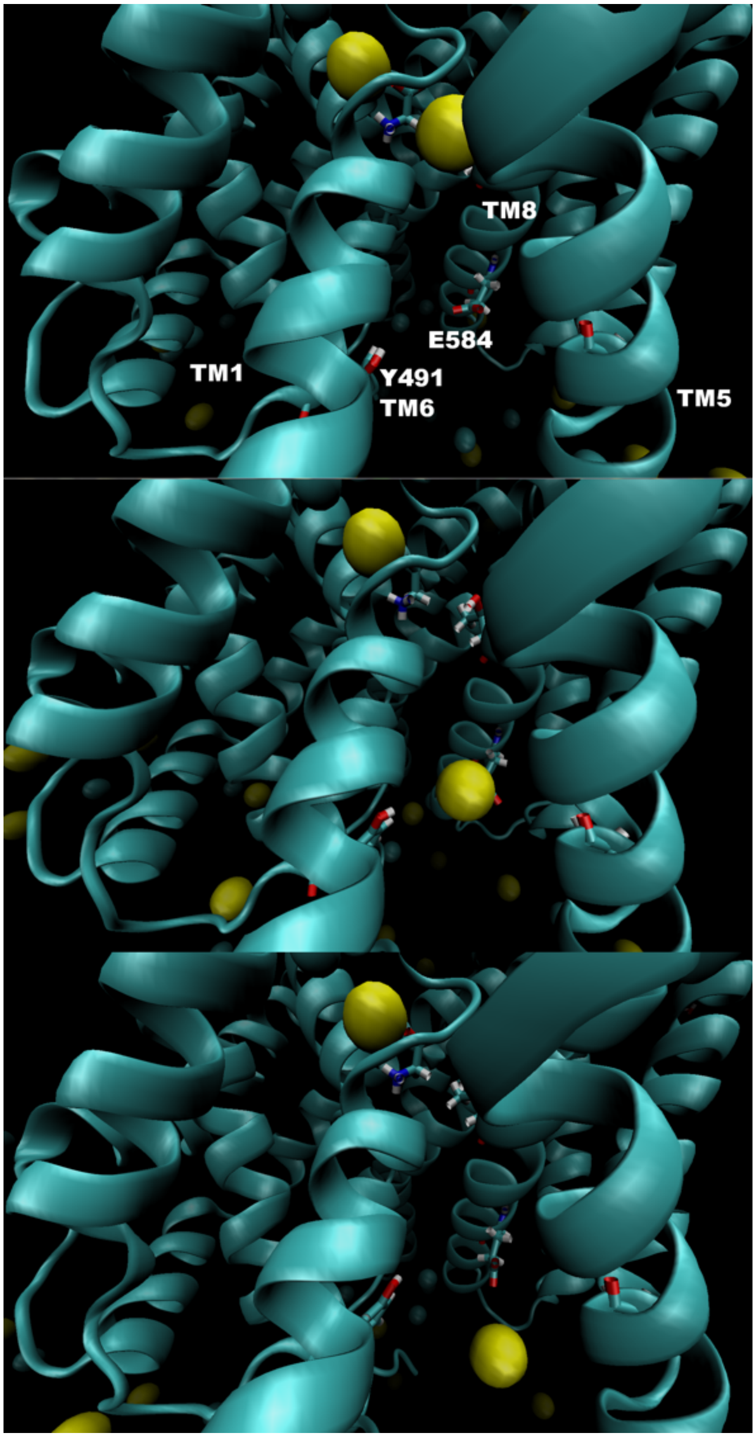

Figure S4: Identified unbinding path of the second $Na^+$ ion in GlyT1.

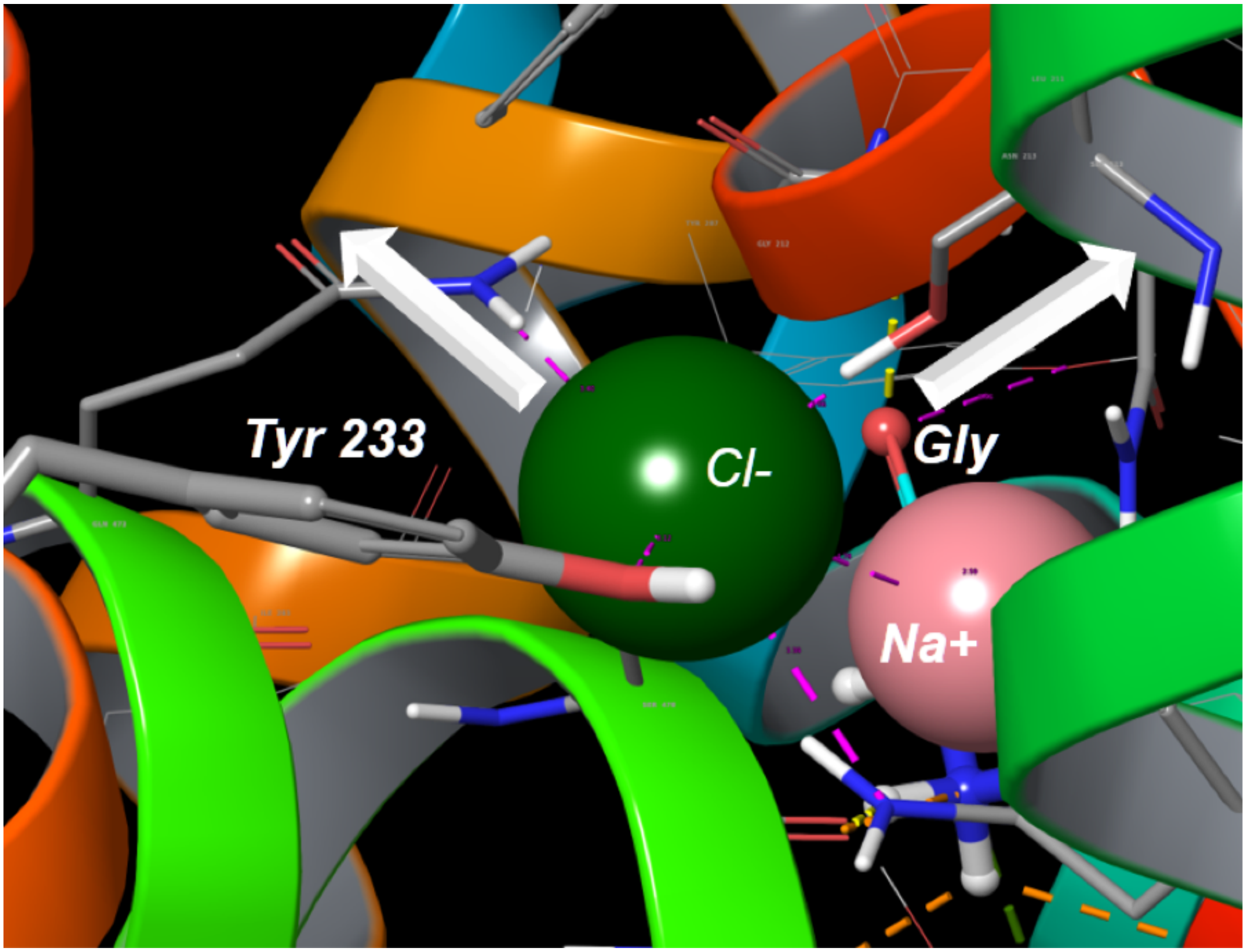

Figure S5: Identified unbinding paths of the second $Cl^-$ ion in GlyT2. With white arrows are shown the unbinding paths of $Cl^-$ (left) and neighboring first sodium ion.

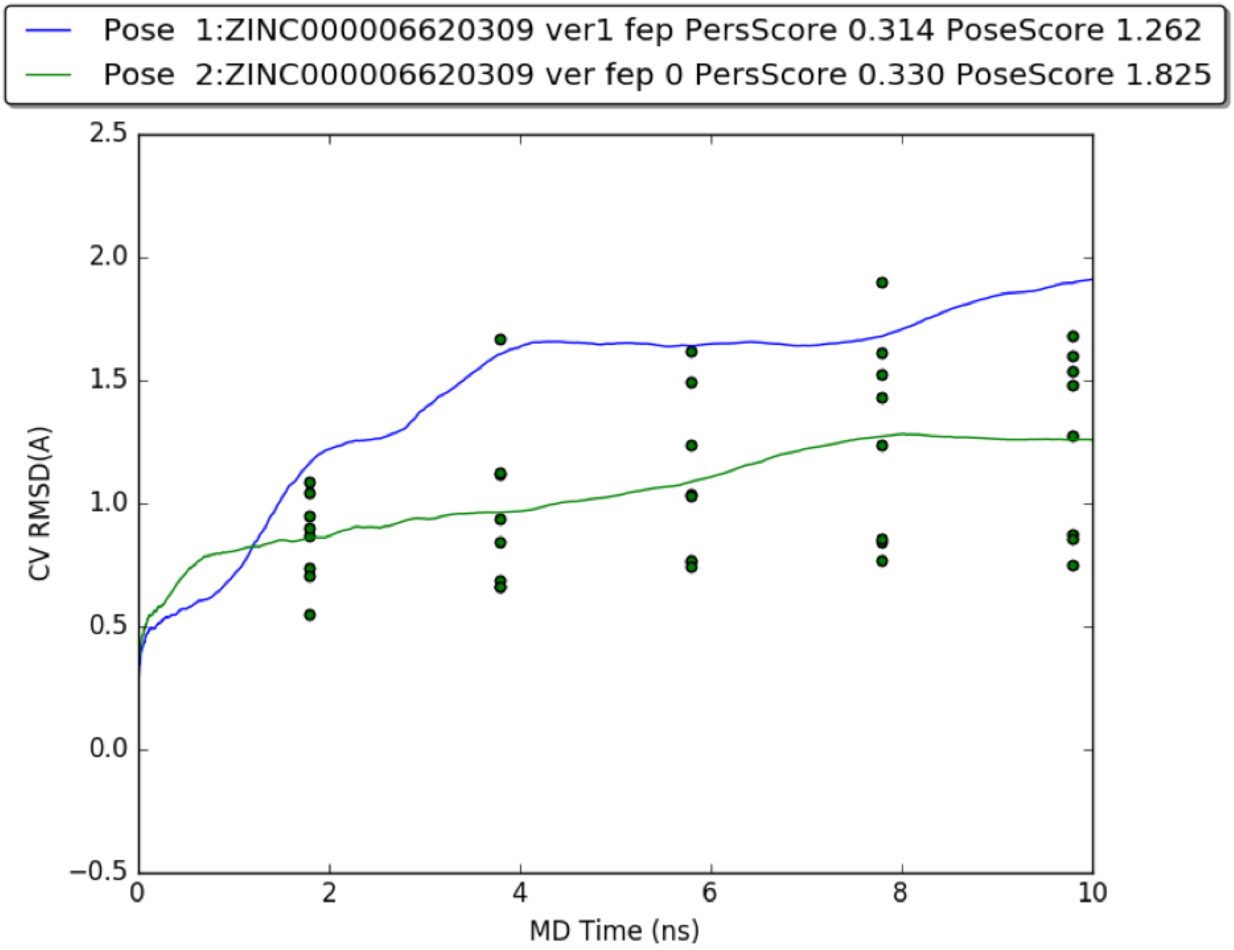

Figure S6: Metadynamics IFD pose prediction results for Lead1. Two docking possess, with rotated by 180 degree aromatic ring, were submit for these calculations and most likely pose was revealed by these runs.

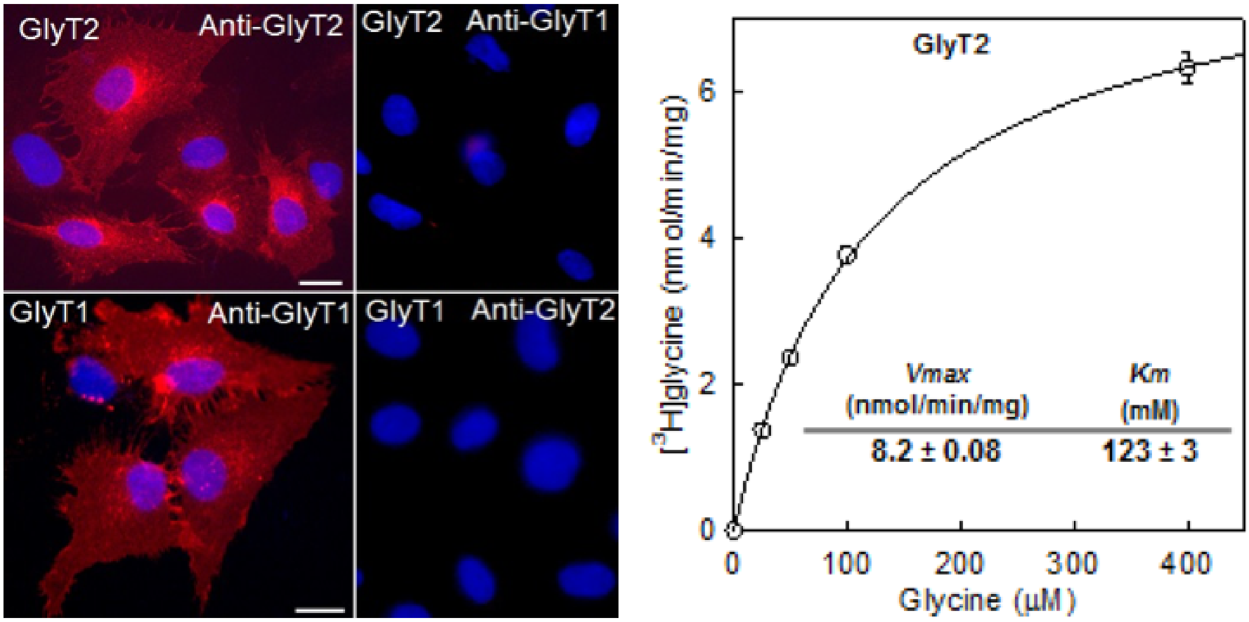

Figure S7: Stable expression of the human GlyT2 and GlyT1b in PAE cells. (A) Cells were fixed with 4% paraformaldehyde followed by staining with anti-GlyT1 or anti-GlyT2 antibodies and CY3-conjugated secondary antibodies. (B) The GlyT2 cells were subjected to uptake assay and the kinetic constants calculated with Sigma Plot 12 software.

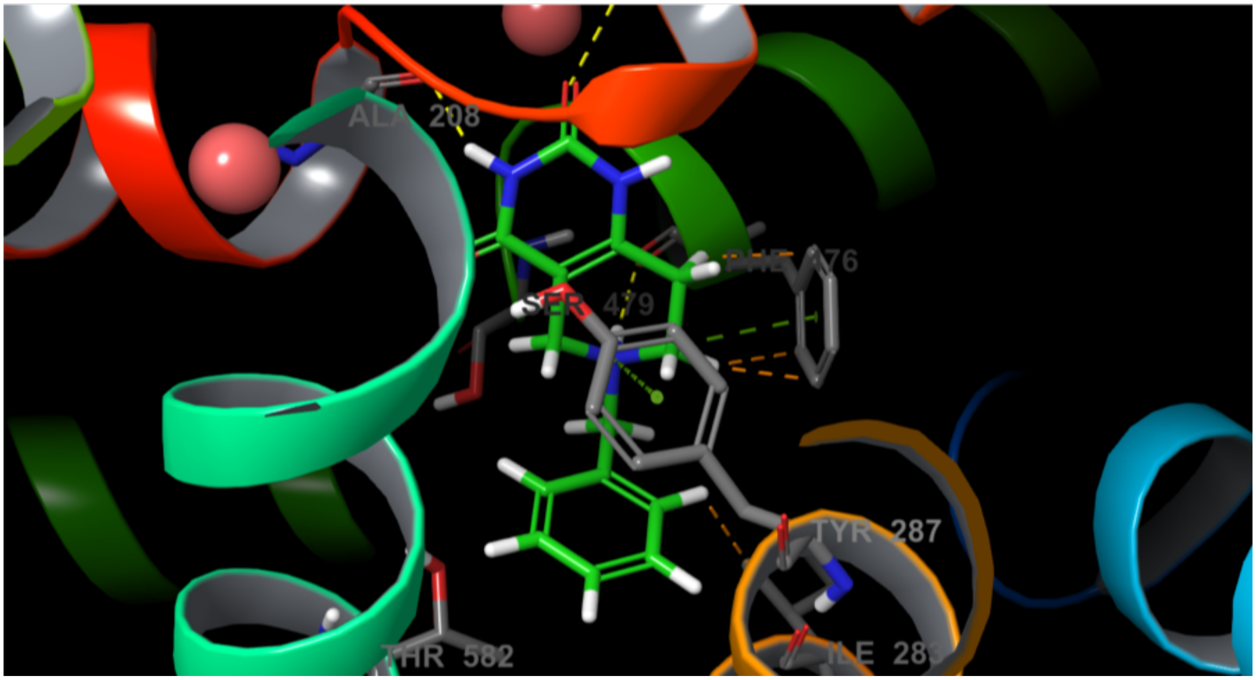

Figure S8: 3D representation of Compound 4-GlyT2 interactions as identified by our MD studies.

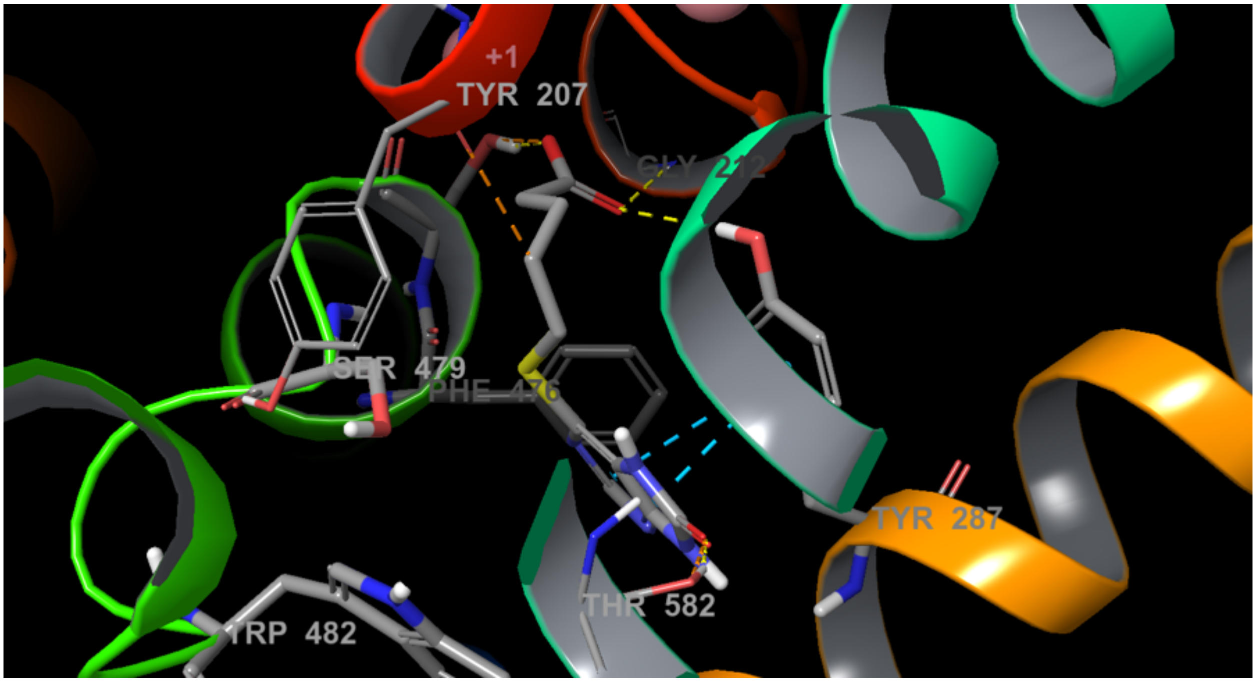

Figure S9: 3D representation of Compound 2-GlyT2 interactions as identified by our MD studies.

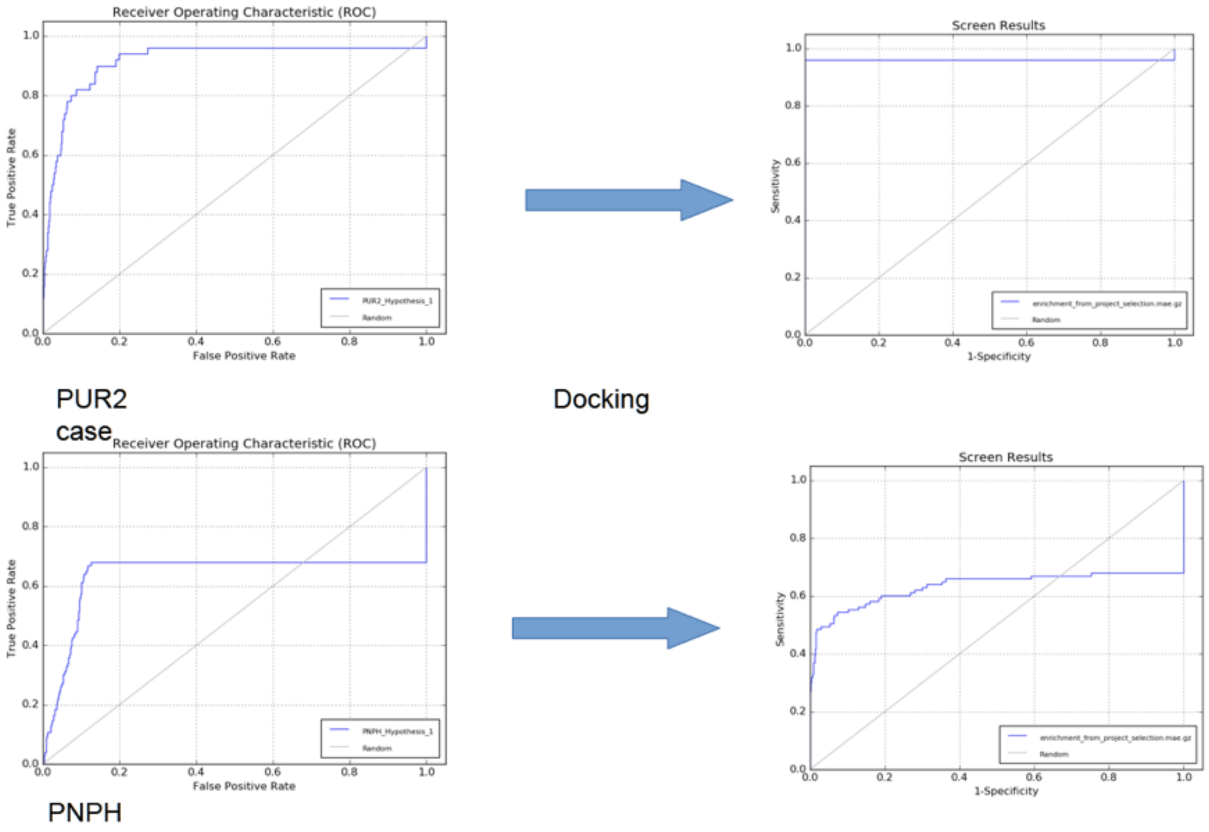

Figure S10: An example of structural based pharmacophore screen results obtained by Phase software (Schrodinger package version 2017-3) on DUD-E. The random selected targets on the figure show an impressive result.

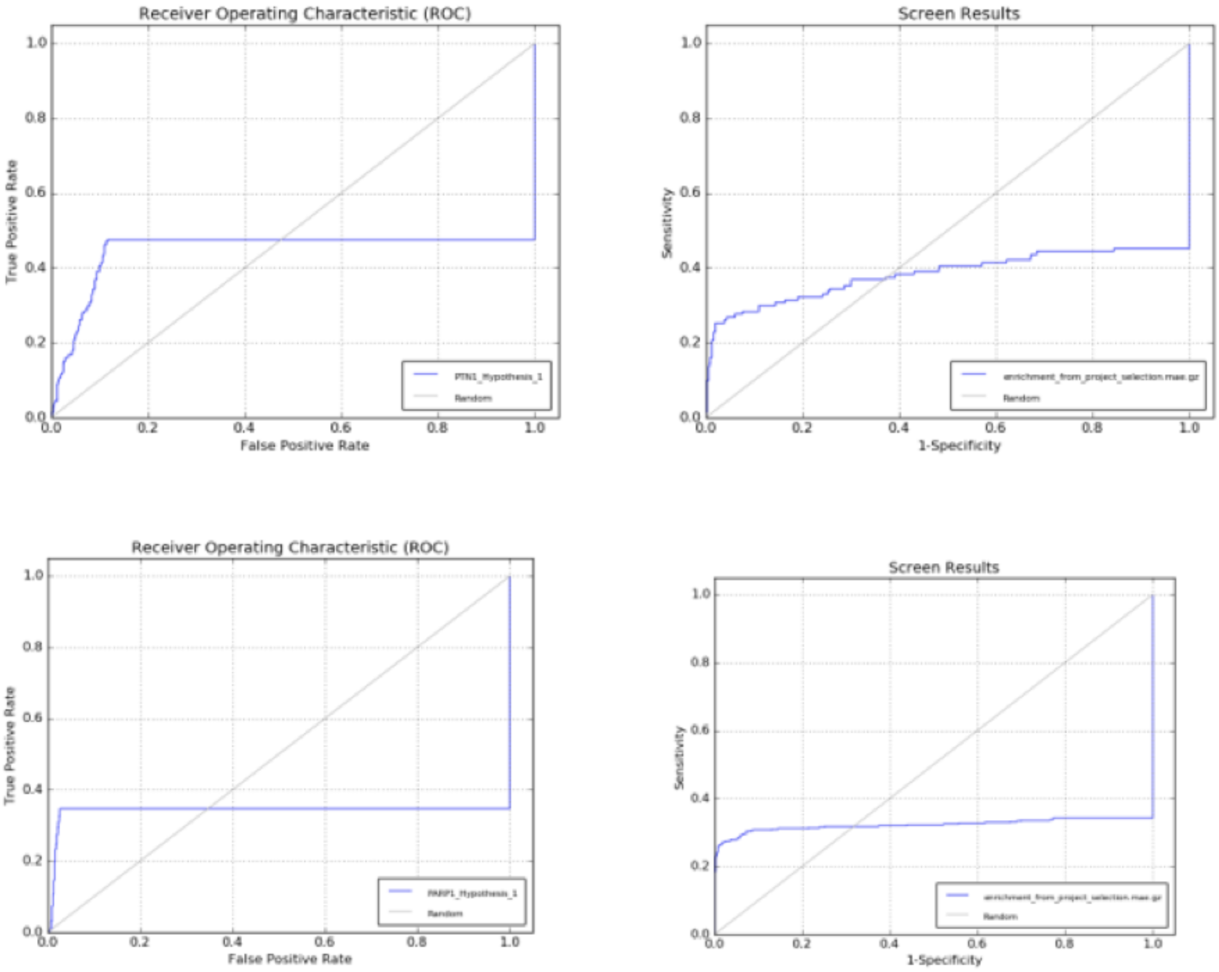

Figure S11: An example of structural based pharmacophore screen results obtained by Phase software (Schrodinger package version 2017-3) on DUD-E. The random selected targets on the figure show a reasonable results.

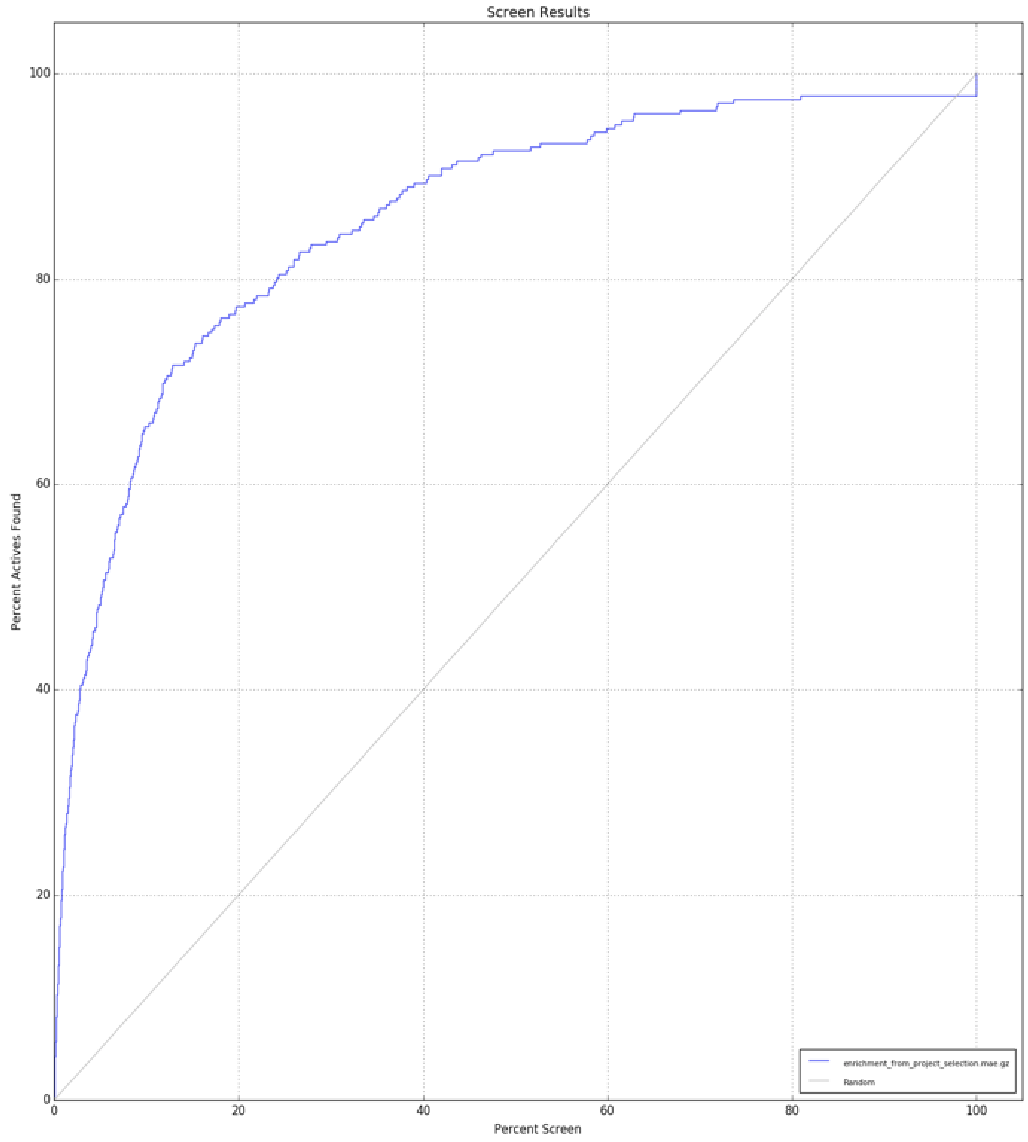

Figure S12: An improved Glide-SP scoring function was detected based on ACE test case

[4] Schrödinger LLC. Schrödinger suite.

[5] Woody Sherman, Tyler Day, Matthew P Jacobson, Richard A Friesner, and Ramy Farid. Novel Procedure for Modeling Ligand/Receptor Induced Fit Effects. *Journal of Medicinal Chemistry*, 49(2):534–553, jan 2006.

[6] Anthony J Clark, Pratyush Tiwary, Ken Borrelli, Shulu Feng, Edward B Miller, Robert Abel, Richard A Friesner, and B J Berne. Prediction of Protein–Ligand Binding Poses via a Combination of Induced Fit Docking and Metadynamics Simulations. *Journal of Chemical Theory and Computation*, 12(6):2990–2998, jun 2016.

[7] Filip Fratev, Thomas Steinbrecher, and Svava Ósk Jónsdóttir. Prediction of Accurate Binding Modes Using Combination of Classical and Accelerated Molecular Dynamics and Free-Energy Perturbation Calculations: An Application to Toxicity Studies. *ACS Omega*, 3(4):4357–4371, apr 2018.

[8] David Case and et. al. AMBER 16, 2016.

[9] Sunhwan Jo, Taehoon Kim, Vidyashankara G. Iyer, and Wonpil Im. CHARMM-GUI: A web-based graphical user interface for CHARMM. *Journal of Computational Chemistry*, 29(11):1859–1865, aug 2008.

[10] Mikhail A Lomize, Irina D Pogozheva, Hyeon Joo, Henry I Mosberg, and Andrei L Lomize. OPM database and PPM web server: resources for positioning of proteins in membranes. *Nucleic acids research*, 40(Database issue):D370–6, jan 2012.

[11] Jean-Paul Ryckaert, Giovanni Ciccotti, and Herman J.C Berendsen. Numerical integration of the cartesian equations of motion of a system with constraints: molecular dynamics of n-alkanes. *Journal of Computational Physics*, 23(3):327–341, mar 1977.

[12] Henrik G. Petersen. Accuracy and efficiency of the particle mesh Ewald method. *The Journal of Chemical Physics*, 103(9):3668–3679, sep 1995.

[13] Edward Harder, Wolfgang Damm, Jon Maple, Chuanjie Wu, Mark Reboul, Jin Yu Xiang, Lingle Wang, Dmitry Lupyan, Markus K. Dahlgren, Jennifer L. Knight, Joseph W. Kaus, David S. Cerutti, Goran Krilov, William L. Jorgensen, Robert Abel, and Richard A. Friesner. OPLS3: A Force Field Providing Broad Coverage of Drug-like

Small Molecules and Proteins. *Journal of Chemical Theory and Computation*, 12(1):281–296, jan 2016.
